# Formerly degenerate seventh zinc finger domain from transcription factor ZNF711 rehabilitated by experimental NMR structure

**DOI:** 10.1101/2024.04.06.588434

**Authors:** Antonio J. Rua, Andrei T. Alexandrescu

## Abstract

Domain Z7 of nuclear transcription factor ZNF711 has the consensus last metal-ligand H23 found in odd-numbered zinc-fingers of this protein replaced by a phenylalanine. Ever since the discovery of ZNF711 it has been thought that Z7 is probably non-functional because of the H23F substitution. The presence of H26 three positions downstream prompted us to examine if this histidine could substitute as the last metal ligand. The Z7 domain adopts a stable tertiary structure upon metal binding. The NMR structure of Zn^2+^-bound Z7 shows the classical ββα-fold of CCHH zinc fingers. Mutagenesis and pH titration experiments indicate that H26 is not involved in metal binding and that Z7 has a tridentate metal-binding site comprised of only residues C3, C6, and H19. By contrast, an F23H mutation that introduces a histidine in the consensus position forms a tetradentate ligand. The structure of the WT Z7 is stable causing restricted ring-flipping of phenyalanines 10 and 23. Dynamics are increased with either the H26A or F23H substitutions and aromatic ring rotation is no longer hindered in the two mutants. The mutations have only small effects on the *K*_d_ values for Zn^2+^ and Co^2+^ and retain the high thermal stability of the WT domain above 80 °C. Like two previously reported designed zinc fingers with the last ligand replaced by water, the WT Z7 domain is catalytically active, hydrolyzing 4-nitophenyl acetate. We discuss the implications of naturally occurring tridentate zinc fingers for cancer mutations and drug targeting of notoriously undruggable transcription factors. Our findings that Z7 can fold with only a subset of three metal ligands suggests the recent view that most everything about protein structure can be predicted through homology modeling might be premature for at least the resilient and versatile zinc-finger motif.

## 1. INTRODUCTION

ZNF711 (also known as CMPX1 and ZNF6), is a zinc-finger (ZNF) nuclear transcription factor required for brain development that occurs on the X-chromosome at position Xq21.1-q21.3 (Kleine-Kohlbrecher *et al*. 2010; Lloyd *et al*. 1991). Mutations in ZNF711 are linked with the intellectual development disorder X-linked 97 (XLID97 also known as MRX97 or IDX97) (Tarpey *et al*. 2009; van der Werf *et al*. 2017; Wang J *et al*. 2022). Male individuals with these mutations are often asymptomatic or have mild to moderate intellectual disability, while female carriers are unaffected (van der Werf *et al*. 2017; Wang J *et al*. 2022). There are some 82 genes on the X-chromosome associated with intellectual disability (Kleine-Kohlbrecher *et al*. 2010). ZNF711 is one of eleven of these proteins containing ZNF domains (Wang J *et al*. 2022). The NIH deemed ZNF711 an understudied protein associated with a rare disease, part of an effort to jumpstart research efforts in this area under the program RFA-TR-22-030.

The ZNF711 transcription factor binds to CpG islands (Rhie *et al*. 2018) that occur in about 70% of human promoters near transcription start sites (Deaton and Bird 2011; Kooy 2022). Genes controlled by promoters with CpG islands can be silenced by methylation of cytosine to 5-methylcytosine. Conversely, unmethylated CpG islands mark chromatin structure competent for transcription. Transcription is further regulated by posttranslational modifications such as acetylation and methylation of histone nucleosome components (Deaton and Bird 2011). KDM5C is a lysine-specific histone demethylase that acts in chromatin remodeling and epigenetic regulation of transcription for a hub of genes involved in neural development (Poeta *et al*. 2019). The ZNF711 transcription factor in concert with the ‘eraser’ histone demethylase PH8F activates transcription of the KDM5C. A second transcription factor ARX also induces KDM5C gene expression and may act as an antagonist of ZNF711, competing with the latter for binding to PH8F (Poeta *et al*. 2019). All four genes *ZNF711*, *KDM5C*, *PH8F*, and *ARX* exhibit mutations connected with X-linked intellectual disability (Kleine-Kohlbrecher *et al*. 2010; Kooy 2022; Poeta *et al*. 2019). Recently, distinct changes in genomic methylation patterns called an “episignature” were identified for individuals afflicted with ZNF711 mutations, the first such example amongst ZNF proteins. This episignature may help assess ZNF711 mutations of uncertain clinical significance or serve to diagnose XLID97 (Wang J *et al*. 2022).

Beyond its role in brain development ZNF711 is part of a family of key regulators of the transcriptome affecting thousands of genes, such that its dysfunction can have roles in cell proliferation and cancer (Ni *et al*. 2020; Rhie *et al*. 2018). To date, roles for ZNF711 have been suggested in breast cancer (Li *et al*. 2020), ovarian serous adenocarcinoma (Kang *et al*. 2015), acute myeloid leukemia (Wang JD *et al*. 2019), and resistance to cisplatin chemotherapy in epithelial ovarian cancer (Wu *et al*. 2021). ZNF711 has also been suggested to play a role in thyroid eye diseases through its role in regulation of the genes involved in that pathology (Hu *et al*. 2022).

ZNF711 belongs to a family of closely related transcription regulators that includes ZFX encoded on the X-chromosome, and ZFY on the Y-chromosome (Johnston *et al*. 1998; Lloyd *et al*. 1991; Ni *et al*. 2020; Valleley *et al*. 1992). The domain organization of the three transcription factors is shown in Figure 1A. All three have an acidic domain at the N-terminus thought to be an activation domain that interacts with other proteins that regulate transcription (PH8F in the case of ZNF711), a nuclear localization signal (NLS), and an array of ZNF domains containing a DNA-binding region (Lloyd *et al*. 1991; Ni *et al*. 2020). The ZNF domains beyond domain Z3 are arranged in even/odd two-finger repeats (labeled blue and green in Fig.1A) that have distinctive sequence signatures (Lloyd *et al*. 1991). It has been proposed based on NMR studies of ZFY domains that the characteristic sequences of the odd and even domains could represent functionally distinct DNA-recognition motifs (Kochoyan, Havel, *et al.* 1991; Kochoyan, Keutmann, *et al.* 1991; Weiss *et al*. 1990), although DNA-binding was not studied explicitly. A recent mutagenesis analysis of domain deletions in ZFX found that the last three ZNF domains 11-13 are necessary and sufficient for DNA binding in transcription regulation (Ni *et al*. 2020). Given that the ZNF domains of ZFX share high sequence similarity with those of ZFY (99%) and ZNF711 (87%), it is expected that that the latter two transcription factors will also bind DNA through their last three ZNF domains. The function of the remaining domains 1-10 is an open question. These could bind proteins rather than DNA, acting as intermolecular protein recognition modules to govern heterooligomeric complex assembly. Alternatively intramolecular interactions between the first ten ZNF domains could serve structural roles in the ZNF711 transcription factor (Brayer and Segal 2008; Iuchi 2001). Another possibility is that the first ten ZNFs are cryptic DNA binding sites that only become activate under distinct conditions or in specific tissues (Bird *et al*. 2003; Ni *et al*. 2020). In this regard, it is worth noting that while the ZFX/ZFY/ZNF711 family of transcription factors have high sequence homology and bind to many of the same promoters (Ni *et al*. 2020), the XLID97 genetic disease mutations are only associated with the ZNF711 transcription factor.

**Figure 1.**
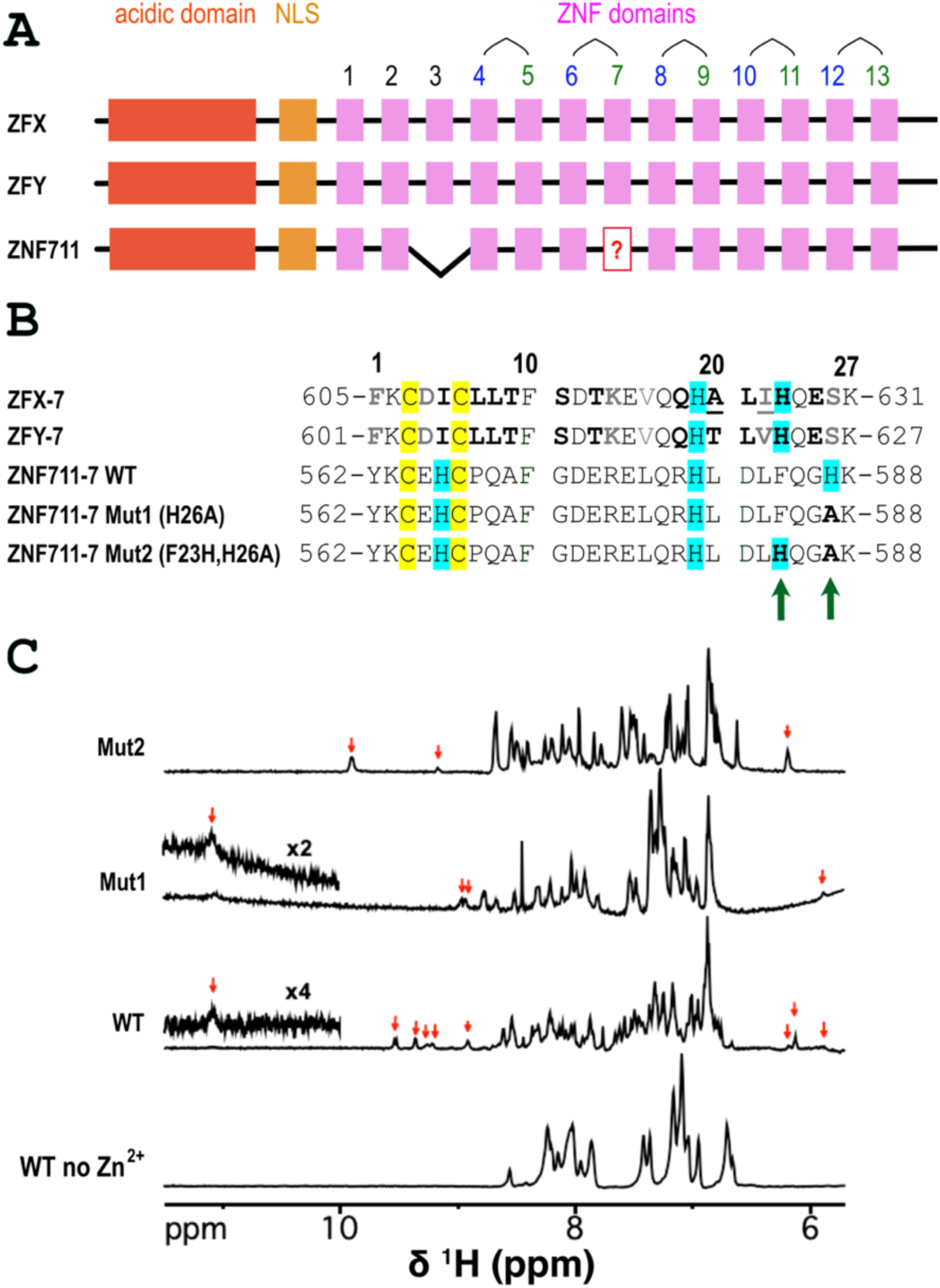
Domain organization, sequence, and NMR spectra of the Z7 domain of ZNF711. **(A)** Domain organization of the closely related transcription factors ZFX, ZF7, and ZNF711. In this paper we use a domain numbering scheme based on that of ZFX. The ZNF domains beyond domain 3 are arranged in couplets, with even and odd-numbered domains having distinct sequence properties. Odd numbered ZNF domains have a three-residue separation between the last two His ligands. Even numbered ZNFs have a four-residue spacing between the last two His residues. In ZNF711, domain Z3 has lost all four Zn^2+^ ligands and is therefore thought to be non-functional. The subject of this paper, domain Z7 was also previously thought to be non-functional due to loss of the fourth zinc ligand H23. **(B)** Sequences of Z7 in the three transcription factors ZFX (605-631), ZFY (601-627), and ZNF711 (562-588), and in two mutant variants of ZNF711-7 used to study Zn^2+^ binding in this work. Putative Zn^2+^ ligands are highlighted yellow for Cys and blue for His. In ZNF711-Z7 the last His ligand at position 23 is replaced by a Phe. The nearby H26 could be the fourth ligand but would give a six-residue spacing between the last two histidines. **(C)** 1D ^1^H NMR spectra of the WT Z7 domain and the two variants Mut1(H26A) and Mut2(F23A,H26A). Red arrows indicate amide and aromatic NMR signals consistent with stable tertiary structure that occur outside the random coil region (7.8 to 8.7 ppm). The insets show a weak NMR signal at 11.09 ppm due to the backbone amide proton of L16, observed in WT and Mut1. Note that the Z7 sequence has no tryptophan sidechains that could account for the 11.09 ppm signal. The 1D NMR data were obtained at 600 MHz on samples at pH 6 and 25 °C.

The UniProt database designates ZNFs as ‘degenerate’ when sequences diverge so much from consensus the resulting domains are thought to have lost the ability to bind Zn^2+^ and are therefore presumed nonfunctional (Aceituno-Valenzuela *et al*. 2020; The_UniProt_Consortium 2023). The folding status of these degenerate ZNFs is currently not easy to predict. We recently reported the NMR structure of the single ZNF domain of ZNF750 (Rua *et al*. 2023). Neither the UniProt database nor AlphaFold were able to correctly establish the folding status of the domain or predict its structure. We corrected examples of false degenerate annotations for three other domains (ZFAT-Z14, ZNF32-Z7, TZAP-Z2) for which there were experimental structures in the Protein Databank (PDB codes 2RV1, 2EPC, 2DLQ), and suggested several domains annotated as degenerate are probably functional CCHC-type ZNFs (Rua *et al*. 2023). More recently we initiated NMR studies of a further five ZNF domains that despite being either annotated as degenerate or not being recognized as ZNFs by UniProt, adopt folded tertiary structure in the presence of Zn^2+^ (A.T.A. & A.J.R., to be published). Incorrect annotation of the folding status of these domains can affect the design of experiments and interpretation of results on ZNF proteins.

For ZNF711, there has been uncertainty since its discovery thirty years ago (Johnston *et al*. 1998; Lloyd *et al*. 1991) as to whether the third and seventh domain of the transcription factor are folded (Kooy 2022; Li *et al*. 2020; Ni *et al*. 2020; van der Werf *et al*. 2017). In this paper we shall use the domain numbering of ZFX for consistency with the other members of this sequence homology family (Figure 1). The segment in ZNF711 corresponding to domain Z3 in ZFX/ZFY (UniProt accession code Q9Y462, residues 448-476) is missing all four of the Zn^2+^-chelating residues. Domain Z7 (UniProt accession code Q9Y462, residues 562-584), is thought to be degenerate since it is missing the final conserved His residue (H23) that completes the tetrahedral Zn^2+^ coordination site of its close homologs ZFX/ZFY (Lloyd *et al*. 1991). However, it has been postulated that H26 three positions downstream in the sequence could substitute as a Zn^2+^ ligand for H23 (Lloyd *et al*. 1991; Valleley *et al*. 1992). Towards the goal of obtaining structural information on rare disease proteins such as ZNF711 beyond gene identification and proteome interaction assessment, we set out to determine the folding state, structure, metal-binding, and dynamic properties of the unconventional Z7 domain.

## 2. RESULTS

### 2.1 ZNF711-Z7 is a genuine ZNF that folds in the presence of Zn^2+^

The sequence of the WT Z7 domain form ZNF711 is shown in Fig. 1B. The consensus last Zn^2+^ ligand for the ZFX/ZFY/ZNF711 family, H23, is missing in the Z7 domain of ZNF711(Lloyd *et al*. 1991). However, H26 three positions downstream in the sequence could substitute in this role for the Z7 domain, or D21 since glutamine or aspartate can sometimes serve as a Zn^2+^ ligand (Kluska *et al*. 2018a). In the absence of Zn^2+^ the NMR spectrum of Z7 is typical of an unfolded polypeptide with all backbone amide hydrogens having chemical shifts in the ‘random coil’ region between 7.8 and 8.7 ppm (Fig. 1C). The NMR spectrum of Z7 experiences large changes in the presence of Zn^2+^, consistent with folding to a stable tertiary structure. While the folded Z7 domain lacks high field shifted methyl resonances upfield of 0.6 ppm, it has several amide proton resonances downfield of 8.9 ppm as well as aromatic resonances downfield of 6.5 ppm (indicated by red arrows) characteristic of folded structure. Aromatic protons in unfolded polypeptides sometimes resonate outside of the random coil region due to ring current shifts induced by residual structure (Mitchell *et al*. 2022; Smith *et al*. 1994). The resonances indicated by red arrows in Fig. 1C are specifically associated with Zn^2+^-binding, and are not seen in the unfolded Z7 domain in the absence of Zn^2+^. The signals from the unfolded and folded domain are in slow exchange on the NMR timescale, typical of ZNFs that usually bind Zn^2+^ with a tight sub-nanomolar dissociation constant (Matousek and Alexandrescu 2004; Rua *et al*. 2023).

### 2.2 The Z7 domain has a classical ZNF ββα fold

NMR assignments for the Z7 domain were obtained using 2D NMR homonuclear (NOESY, TOCSY) and heteronuclear (^1^H-^13^C HSQC and ^1^H-^15^N sofast-HMQC) experiments as described in the Methods section. The quality of the data is illustrated by the fingerprint Hα−Cα region from the ^1^H-^13^C HSQC spectrum recorded at natural ^13^C isotopic abundance (Fig. 2A). The Hα chemical shifts range between 4.9 ppm and 3.9 - 2.9 ppm for some of the residues in the α-helix. The bulk of the NMR assignments were done at a temperature of 25 °C. During the start of structure determination, we noticed that the amide proton of L16, which is barely visible at 11.09 ppm (Fig. 1C) becomes stronger at 10 °C (Fig. 2B) and therefore collected all the NOESY data for structure determination at the lower temperature. NMR assignments were readily transferrable between the two temperatures, and the data at 10 °C allowed us to complete assignments and define the role of L16 in the structure.

**Figure 2.**
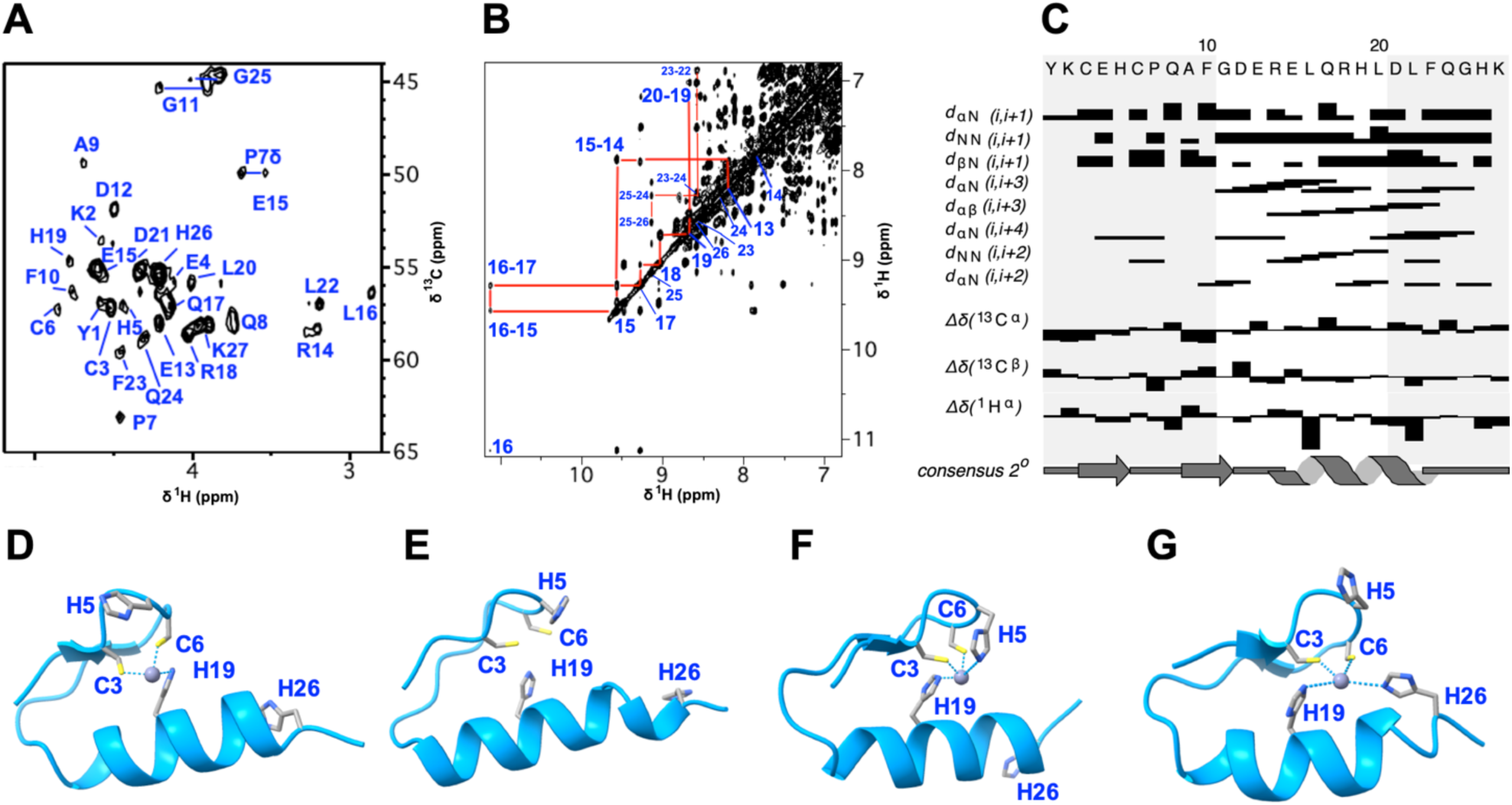
NMR spectra and structure of Z7 WT. **(A)** Assigned ^13^C-HSQC obtained at pH 6.2, 25 °C for a sample of Z7 at natural isotopic abundance. **(B)** Portion of the 250 ms 2D NOESY spectrum at 10 °C showing an α**-**helix d_NN_ assignment walk from residues 13 to 26. **(C)** Wüthrich diagram of short-range NOEs and chemical shift differences sensitive to secondary structure. **(D)** NMR structure of the Z7 conformer closest to the ensemble average (model 1). **(E)** Colabfold prediction of the Z7 structure. **(F)** Representative of about 25% of the experimental Z7 NMR conformers, where H5 comes within metal bonding distance of Zn^2+^. **(G)** NMR structure of Z7 calculated with a Zn^2+^-bonding restraint to H26. Introduction of this artificial restraint caused no distance or dihedral violations and gave very similar energies to the structures calculated without it.

Fig. 2C summarizes the NOE and chemical shift patterns pertaining to the secondary structure of the Z7 domain. The data are consistent with a ββα secondary structure, typical of the classical ZNF folding motif (Klug 2010; Lee *et al*. 1989; Neuhaus 2022; Padjasek *et al*. 2020). Fig. 2D shows the NMR structure of Z7 closest to the ensemble average, calculated from the restraints summarized in Table 1. The structure has the classical ZNF fold, giving a backbone RMSD of 1.6 Å to the prototypical middle domain from the 1.6 Å-resolution X-ray structure of ZIF268 ((Elrod-Erickson *et al*. 1998), PDB code 1A1H).

**Table 1.** Statistics for the 20 best NMR structures of the Z7 domain from ZNF711.

|  |  |  |
| --- | --- | --- |
| NMR restraints (total) | 240 |  |
| Distance (total) | 185 |  |
| Intraresidue NOE ($\vert i-j\vert = 0$) | 47 | |
| Sequential NOE ($\vert i-j\vert = 1$) | 57 | |
| Medium-range NOE ($1 < \vert i-j\vert < 5$) | 32 | |
| Long-range NOE ($\vert i-j\vert \geq 5$) | 18 | |
| Hydrogen bond <sup>a</sup> | 26 |  |
| Restraints to Zn<sup>2+</sup> atom <sup>b</sup> | 5 |  |
| Dihedral ($\phi$ 21, $\psi$ 21, $\chi_1$ 13) | 55 | |
| *Restraint violations* |  |  |
| > 0.5 Å | 0 |  |
| > 5° | 0 |  |
| *Residual restraint violations* |  |  |
| Distance (Å) | 0.0811 ± 0.0037 <sup>c</sup> |  |
| Dihedral (°) | 0.77 ± 0.19 |  |
| *RMS deviations from ideal geometry* |  |  |
| Bonds (Å) | 0.00258 ± 0.00024 |  |
| Angles (°) | 0.47 ± 0.03 |  |
| Improper torsions (°) | 0.30 ± 0.02 |  |
| *Ramachandran plot PROCHECK statistics for ordered residues* |  |  |
| Residues in most favored regions | 87.9 % |  |
| Residues in allowed regions | 12.1 % |  |
| Residues in generously allowed regions | 0.0 % |  |
| Residues in disallowed regions | 0.0 % |  |
| *Coordinate rms deviations (Å) <sup>d</sup>* |  |  |
|  | <u>Backbone (Cα,N,C')</u> | <u>All</u> |
| NMR ensemble to mean | 0.83 ± 0.19 | 1.55 ± 0.24 |
| NMR to Colabfold | 1.64 ± 0.23 | 2.72 ± 0.34 |
| NMR ensemble to PDB 1AH1 135-156 <sup>e</sup> | 1.56 ± 0.21 | N.A. |

<sup>a</sup>Two distance restraints (1.5-2.5 Å for NH-O and 2.5-3.5 Å for N-O) were included for each of the 13 H-bond restraints to enforce H-bond linearity.
<sup>b</sup>Two restraints per Cys (from S $\gamma$ and Cβ to maintain linearity) and one restraint per His (Nδ1 or Nε2)
<sup>c</sup>All values are given as mean ± standard deviation.
<sup>d</sup>Alignments were done for the entire structure using the PyMol program with cycles=0. No atoms were rejected for the calculation of RMSD values.
<sup>e</sup>PDB 1AH1 is a 1.6 Å resolution X-ray structure of a ZIF268 variant in complex with DNA. Residues 135-152 correspond to the middle domain and are used here as a benchmark for a 'generic' zinc finger.

### 2.3 The WT Z7 domain has only three Zn^2+^-ligating residues

The final Z7 NMR structure has only three Zn^2+^ ligands C3, C6, and H19 (Fig 2D), however establishing that the domain has an unusual tridentate coordination site required additional NMR and mutagenesis data as described below. Fig 2E shows a model of the Z7 domain calculated with the ColabFold program (Mirdita *et al*. 2022). Note that the AlphaFold algorithm (Jumper *et al*. 2021) used by ColabFold does not consider cofactors, so the Zn^2+^ ion is not modeled in the structure. In contrast to the Z* domain from ZNF750 we recently described where AlphaFold models the structure incorrectly (Rua *et al*. 2023), the ColabFold model for Z7 is very close to the NMR structure giving a backbone RMSD of 1.6 Å (Table 1). Like in the NMR structure, H26 in the ColabFold model is too far to be involved in the Zn^2+^ coordination site (Fig. 2E).

In about 25% of the NMR conformers, the Nε2 or Nδ1 atoms of H5 come within 3.5 Å of the Zn^2+^ ion suggesting that this residue may participate in metal binding. The geometry deviates from a tetrahedral Zn^2+^ coordination sphere, however, as illustrated by a representative conformer in Fig. 2F. To explore if the last histidine, H26 could participate in Zn^2+^ binding we introduced a restraint from the Nε2 or Nδ1 atoms of the residue to the metal, and retained the three previously included restraints from C3, C6, and H19. Remarkably, inclusion of the H26 restraint results in a structural model with a tetrahedral Zn^2+^ coordination site (Fig. 2G), introduces no NMR restraint violations, and has an X-plor energy function score nearly identical to that obtained with just three restraints to the metal. The only difference between this assumed model and the correct NMR structure for Z7 is that the C-terminus of the α-helix is slightly bent and distorted when the metal restraint from H26 is included (Fig. 2G). While C6 and H19 show long-range NOEs to residues across the Zn^2+^ binding site including between C6-F23 and H19-F10, H26 only has two sequential NOEs to G25 and K27. Moreover, H26 has a chemical shift-predicted *S*^2^ order parameter of 0.6, the second lowest besides the C-terminus K27, suggesting the residue is flexible. Metal bonding restraints are typically assumed in NMR structure calculations, but the current example illustrates that the NOESY-derived NMR restraints alone cannot distinguish between a tridentate and tetradentate Zn^2+^ coordination site for the Z7 domain.

To better ascertain which histidine residues are involved in Zn^2+^ coordination, we obtained pH titration data. We reasoned that non-bonded histidines should titrate according to their microscopic p*K*_a_ values, whereas metal-bonded histidines should have restricted protonation. These latter sites would either experience no chemical shift changes with pH, or very perturbed p*K*_a_ values. The approach was previously used to distinguish histidines participating in Zn^2+^ coordination by the Nos99 ZNF fragment of the Nanos protein (Curtis *et al*. 1997). For the WT Z7 domain we see the signals of H5 and H26 titrate with pH (Fig, 3A,D) while H19 is invariant with pH. This indicates that in the WT Z7 domain H19 coordinates Zn^2+^, whereas H5 and H26 do not. Residue H5 has a considerably lowered p*K*_a_ of 6.0 suggesting it is in an environment that resists protonation. This could be consistent with H5 stabilizing the bound Zn^2+^ as suggested by the proximity of H5 to the metal in 25% of the NMR structures. Significant broadening of the Hε1 signal of H5 is suggestive of a conformational exchange process on the µs-ms timescale. The fact that H5 titrates with pH, however, indicates that this histidine is not directly bonded to Zn^2+^ but may be participating in the second coordination sphere for the metal, possibly through other groups in the polypeptide or water.

As a further check of the involvement of H26 in Zn^2+^ binding we synthesized the H26A variant, abbreviated Mut1. The Mut1 variant proved much more difficult to work with than WT as it had poor solubility, a CD spectrum closer to the unfolded peptide in the absence of Zn^2+^ than the other two variants (Fig, 4A), and a reduced NMR chemical shift dispersion (Fig. 1C). We initially missed that the Mut1 variant folds in the presence of Zn^2+^ as we only observed only two weak amide protons outside the random coil range near 8.9 ppm. More careful inspection of the 1D NMR spectrum shows a peak at 5.87 ppm due to the Hε aromatic ring hydrogens of F10 and the weak signal at 11.09 ppm due to the backbone amide proton of L16 that is also seen in the WT (Fig. 1C). These signals are incompatible with an unfolded structure and show that Mut1 adopts a stable tertiary structure upon binding Zn^2+^ even when H26 is substituted by an alanine. Therefore, H26 is dispensable for Zn^2+^ binding and folding. A pH titration experiment for Mut1 shows that H19 is invariant to pH changes due to bonding to Zn^2+^ as it is in WT, and that H5 titrates with a lowered p*K*_a_ of 6.0 similar to that in the WT (Fig, 3B,E).

In summary, the pH titration and mutagenesis data indicate that of the three histidines in the sequence of the WT Z7 domain only H19 is bonded to Zn^2+^, forming a tridentate binding site for the metal together with C3 and C6.

### 2.4 A fourth ligand introduced at the H23 consensus position is bonded to Zn^2+^

Since the WT Z7 domain can bind Zn^2+^ with only three amino acids, we wondered what would happen if we replaced the non-conserved F23 residue in this domain with the histidine that occurs at position 23 in all the other viable odd-numbered ZNF domains in the ZNF711/ZFX/ZFY family (Lloyd *et al*. 1991; Ni *et al*. 2020). Would the new H23 introduced at the consensus position participate in Zn^2+^ binding? To this end, we synthesized the double mutant F23H+H26A, abbreviated as Mut2 (Fig. 1B). In analogy with Mut1, the H26A mutation removes the possibility that H26 could participate in Zn^2+^ ligation. The second F23H mutation in Mut2 introduces the consensus histidine that could complete a tetradentate Zn^2+^ binding site, starting from the tridentate platform provided by WT residues C3, C6, and H19. Mut2 is the best behaved amongst the three variants we looked at. It gives 1D NMR spectra characteristic of folded structure (Fig. 1C). Although we did not observe the 11.09 ppm resonance corresponding to L16 HN in Mut2, a new signal at 9.90 ppm could correspond to this proton. The pH titration data on Zn^2+^-bound Mut2 (Fig. 1C) shows that two histidines assigned to H19 and H23 have chemical shifts invariant with pH, and that H5 titrates as it did in WT and Mut1. Based on these data we conclude that Mut2 has a standard tetradentate Zn^2+^ coordination stie comprised of C3, C6, H19, and H23. The p*K*_a_ of H5 in Mut2 is slightly raised towards normal values in Mut2 compared to WT and Mut1 (Fig. 3F). Perhaps when H23 completes the tetradentate metal-coordination site in Mut2, H5 makes less of a contribution to Zn^2+^ binding.

**Figure 3.**
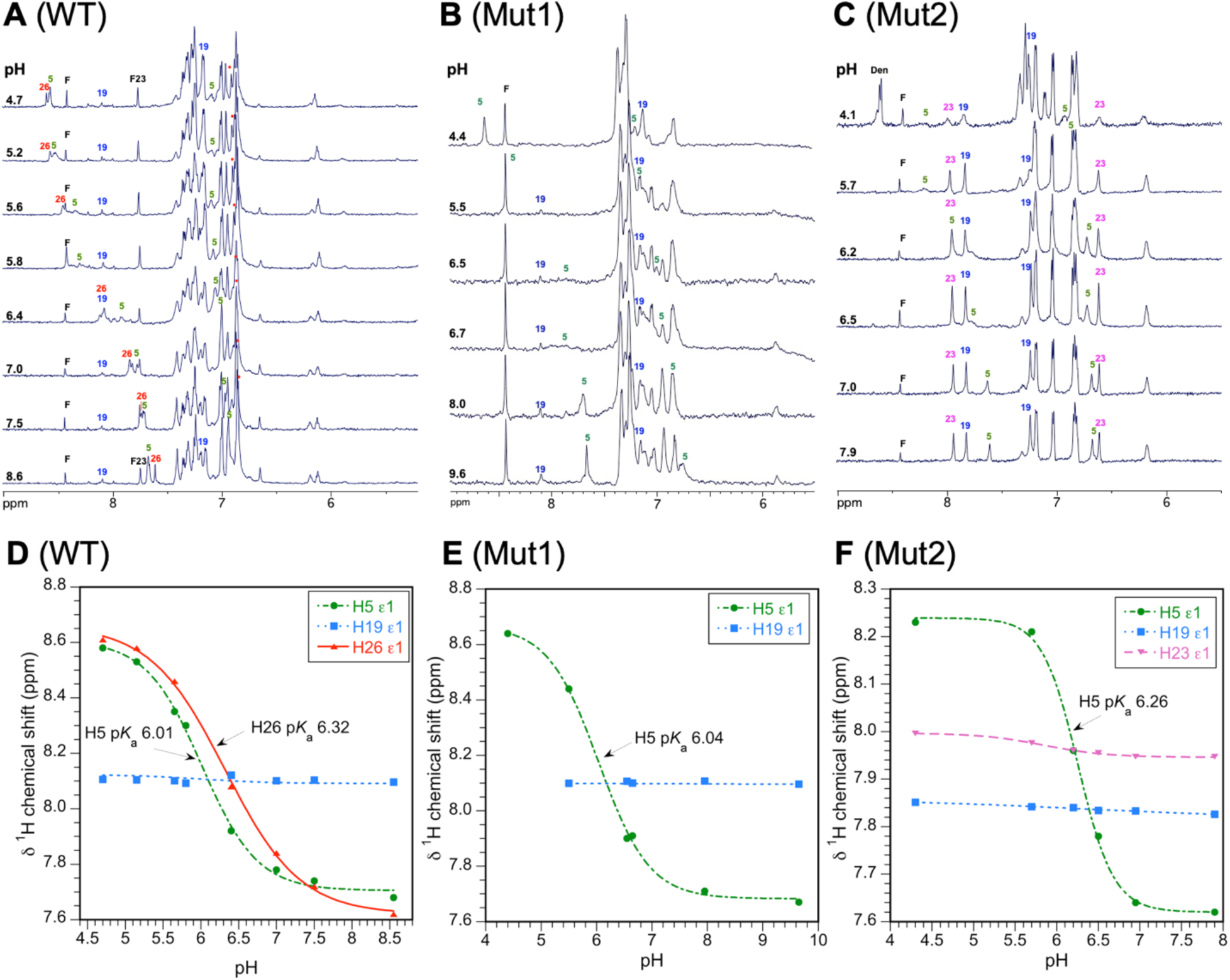
pH titrations used to identify histidines bonded to Zn^2+^ in the Z7 domain. Downfield regions of 500 MHz NMR spectra in D_2_O as a function of pH recorded at a temperature of 25 °C for (**A**) WT, (**B**) Mut1, and (**C**) Mut2. Labeled resonances downfield of 7.5 ppm are due to His Hε1 resonances, those upfield of 7.5 ppm are due to His Hδ2 resonances (labeled with dots of the same colors as the Hε1 resonances in cases of spectral crowding). F is an impurity (formate). (**D-F**) NMR pH titration data for each of the His Hε1 resonances in WT and mutants. Resonances of histidines bonded to Zn^2+^ do not shift. The p*K*a values for the other non-ligating histidines are given in the figure.

### 2.5 Dissociation constants for Zn^2+^ and Co^2+^ are similar for WT and the metal-coordination mutants

We next wanted to see how the WT, Mut1, and Mut2 variants compared in binding of Zn^2+^ and Co^2+^ ions. Fig. 4A compares CD spectra of the Zn^2+^-bound states of the three variants to those of the unfolded WT domain in the absence of Zn^2+^. WT and Mut2 show more negative ellipticity at 208 and 220 nm in their Zn^2+^-bound states, characteristic of an increase in α-helical structure compared to the unfolded state. The effects are small but comparable to those for the folding of other tridentate ZNFs reported in the literature (Kluska *et al*. 2018a). As previously mentioned, the CD spectrum of Mut1 is close to that of the unfolded peptide (Fig. 4A) but NMR shows resonances consistent with stable tertiary structure (Fig. 1C). We used CD spectroscopy to obtain information on Zn^2+^ avidity in a competition assay with EGTA (Rua *et al*. 2023). For Mut2 we used the CD signal at 222 nm to follow the Zn^2+^ titration (Fig 4B), whereas 220 nm was used for WT, and 214 nm for Mut1. The Zn^2+^ binding curves for the three variants are shown in Fig. 4C, and K_d_ values calculated from fits of the data to the Hill equation are given in Table 1. The WT domain has the lowest *K*_d_ for Zn^2+^ at 1.4 x10^-12^ M. The Mut2 variant with a tetradentate ligating site has a *K*_d_ 2-fold higher than WT. The tridentate Mut1 variant has a shallower binding curve suggestive of lowered binding cooperativity, and a K_d_ five-fold larger than WT. All three variants maintain remarkably close K_d_ values in the pM range for Zn^2+^-binding.

**Figure 4.**
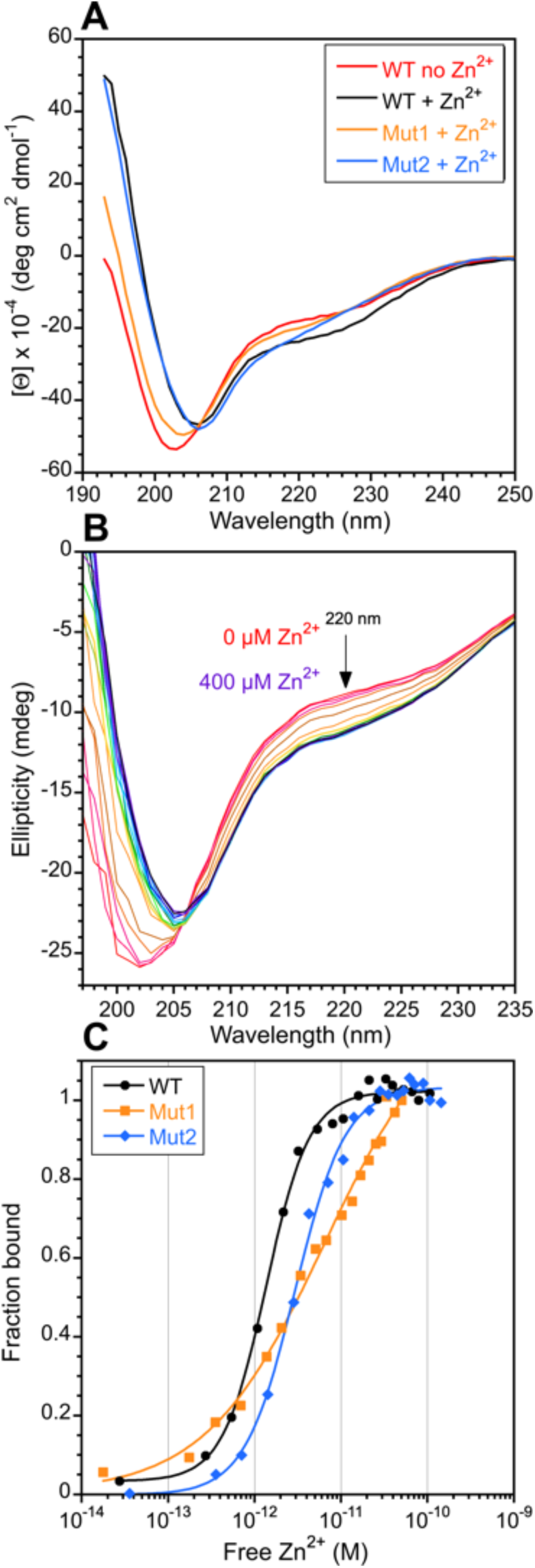
Analysis of Zn^2+^ binding by CD spectroscopy. **(A)** Spectra of WT in the absence of Zn^2+^, and WT and mutants in the presence of 1.2:1 Zn^2+^. **(B)** Titration of 75 μM Mut2 in the presence of 18 mM of the competitor EGTA at 22 °C and pH 7. **(C)** Zn^2+^ binding curves for WT, Mut1, and Mut2. Dissociation constants calculated from the binding curve fits are given in Table 2.

**Table 2.** Apparent dissociation constants (M) for Zn^2+^ and Co^2+^ binding.

| <u>Zinc finger</u> | <u>K<sub>d</sub></u><sup>Zn</sup> | <u>K<sub>d</sub></u><sup>Co</sup> |
| --- | --- | --- |
| WT <sup>a</sup> | 1.36 (± 0.07) x 10<sup>-12</sup> | 2.46 (± 0.26) x 10<sup>-5</sup> |
| Mut1 | 6.50 (± 0.20) x 10<sup>-12</sup> | 2.28 (± 0.35) x 10<sup>-5</sup> |
| Mut2 | 2.94 (± 0.17) x 10<sup>-12</sup> | 4.60 (± 0.23) x 10<sup>-5</sup> |

<sup>a</sup>Uncertainties in parentheses are the standard errors obtained from non-linear least squares fits of the metal binding data to the Hill equation.

We next looked at Co^2+^ binding since UV-vis spectrophotometry of the transition metal is highly sensitive to the environment of the coordination site (Fig 5). The ligand-to-metal charge transfer (LMCT) band depends on the number of thiolate groups bound to the Co^2+^ ion, with the extinction coefficient at 320 nm averaging 900-1200 M^-1^cm^-1^ per Cys-S^-^-Co^2+^ bond (Nomura and Sugiura 2004a). At saturating Co^2+^ concentrations the ε_320 nm_ values are WT 2150, Mut1 2360, and Mut2 2650 M^-1^ cm^-1^, consistent with each variant using two cysteines (C3 and C6) to bind Co^2+^. The d-d transition bands between about 550-750 nm (Fig. 5A inset, Fig. 5B) are sensitive to the coordinating atom types (Krizek *et al*. 1993; Sivo *et al*. 2017) and to the geometry of the Co^2+^ coordination site (Kluska *et al*. 2018b). Tetrahedral coordination results in extinctions coefficients > 300 M^-1^ cm^-1^, pentacoordinate environments in the range 50-300 M^-1^ cm^-1^, and octahedral or hexacoordinate geometry give the lowest extinction coefficients <50 M^-1^ cm^-1^ (Kluska *et al*. 2018a). All three variants show absorption maxima at 568 and 634 nm that are typical of a tetrahedral coordination site in a CCHH-type ZNF (Krizek *et al*. 1993; Sivo *et al*. 2017). The Mut2 mutant which has a tetradentate ligating set has the most intense d-d absorption profile with an ε_634 nm_ maximum of 806 M^-1^ cm^-1^. The d-d absorption profiles for the tridentate WT (ε_634 nm_ = 436 M^-1^ cm^-1^) and Mut1 (e_634 nm_ = 326 M^-1^ cm^-1^) variants are broader and weaker than that of Mut2 but the overall shape is retained (Fig. 5B) and the extinction coefficients are in a range consistent with tetrahedral coordination. The results are reminiscent of a designed CCHH-type consensus ZNF called CP-1 (Krizek *et al*. 1991) where truncation of the last four amino acids led to a broadening and a decreased intensity for the d-d portion of the electronic absorption spectrum due to replacement of the final Co^2+^ ligand H24 by a water molecule (Merkle *et al*. 1991).

**Figure 5.**
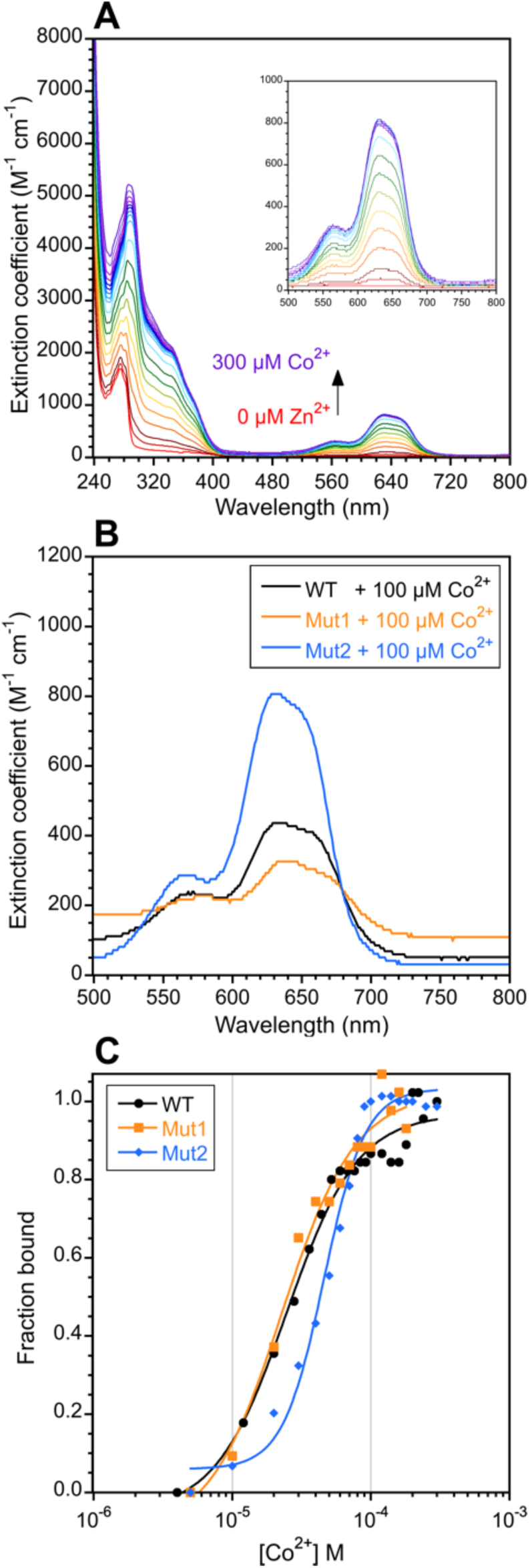
Co^2+^ binding experiments monitored by UV-Vis spectroscopy. **(A)** Co^2+^ titration for Mut2. The inset shows an expansion of the region from 500 to 800 nm. **(B)** Comparison of the d-d transition region of the spectra for 100 µM samples of WT, Mut1, and Mut2 in the presence of 100 µM Zn^2+^. **(C)** Co^2+^ binding curves. All data were obtained at 25 °C. Dissociation constants calculated from the fits are given in Table 2.

Figure 5C shows binding curves obtained from the increase in the ε_634 nm_ absorption maximum with increasing Co^2+^ concentration, and the associated *K*_d_ values for Co2+ obtained are given in Table 2. WT and Mut1 have indistinguishable *K*_d_ values while that of Mut2 is only two-fold higher.

### 2.6 The absence of a fourth Zn^2+^ ligand imparts hydrolytic activity on WT Z7

A characteristic of ZNF domains that have one of the polypeptide ligands replaced by water is a change from a purely structural domain to one that becomes a catalyst (Besold *et al*. 2016; Kluska *et al*. 2018b). Hydrolytic activity has been previously reported for two designed ZNFs: a mutant of the aforementioned CP-1 domain in which the CCHH ligating set was changed to CAHH (Besold *et al*. 2016; Kluska *et al*. 2018b), and several mutants replacing Zn^2+^-coordinating residues by alanine in a ZNF domain from the transcription factor Sp1 (Nomura and Sugiura 2004a). We wondered if the naturally occurring WT tridentate Z7 domain from ZNF711 might also have hydrolytic activity. Figure 6 demonstrates that the Z7 domain hydrolyzes the substrate 4-NA to the chromogenic product 4-NP that can be quantified by absorbance at 400 nm (Fig. 6 A, B). It is interesting to note that the 4-NA substrate is typically used to assay the activity of carbonic anhydrase (Anderson *et al*. 1994; Tu *et al*. 1986), an enzyme with a somewhat similar active site consisting of a Zn^2+^ ion bound by three histidines and a catalytic water molecule in a tetrahedral coordination sphere (Eriksson *et al*. 1988).

**Figure 6.**
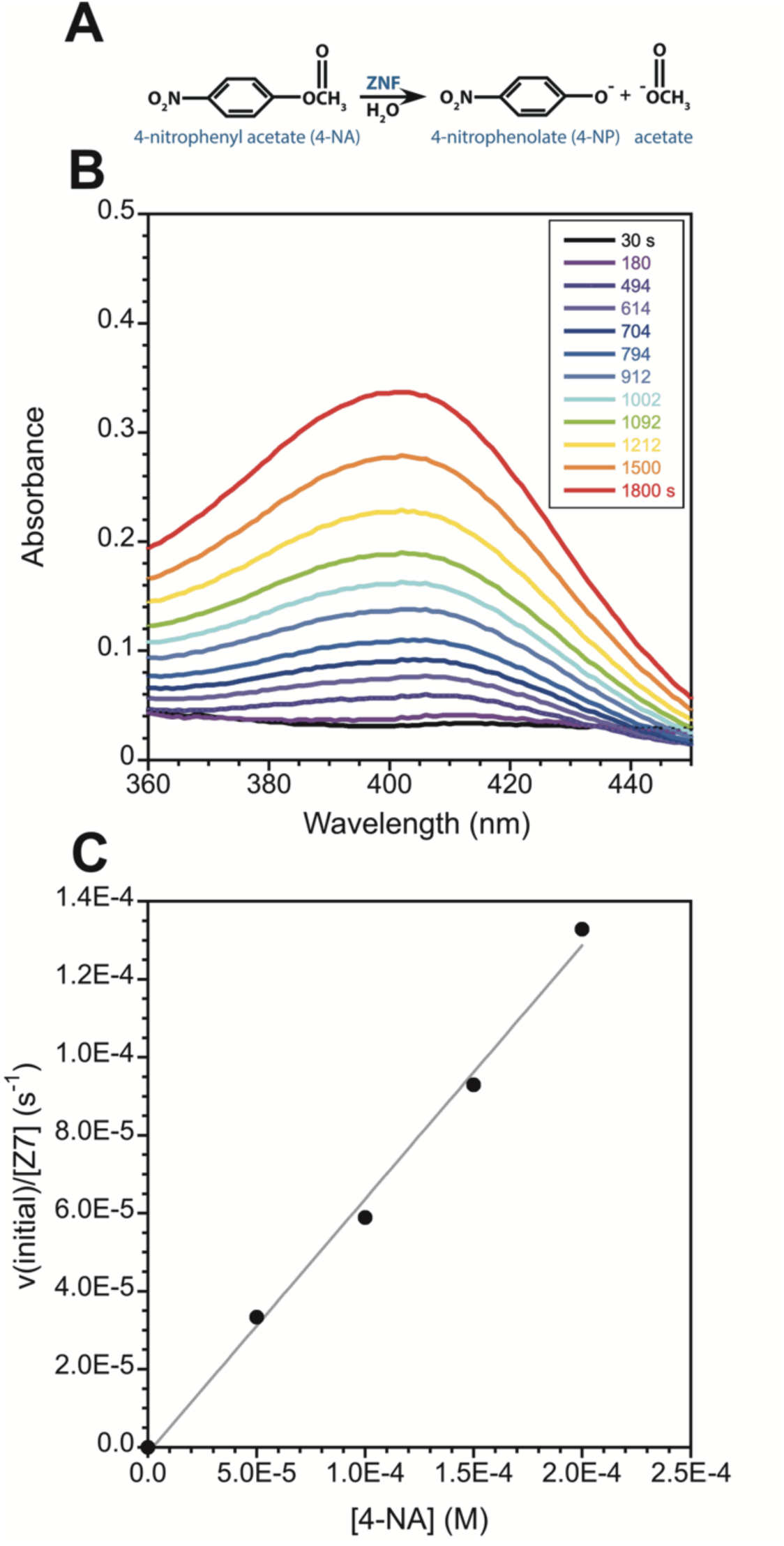
The WT Z7 domain has hydrolytic activity. **(A)** Scheme of the hydrolysis of 4-NA to 4-NP catalyzed by the Zn^2+^-bound tridentate WT Z7 domain. (**B**) Increase in absorbance due to 4-NP production as a function of time. For actual kinetic experiments we monitored the reaction at a single wavelength (ε_400 nm_ = 12,800 M^-1^cm^-1^ for the 4-NP product). (**C**) Plot of initial reaction velocity as a function of 4-NA concentration. The slope gives the second order rate constant k” = 0.65 ± 0.03 M^-1^ s^-1^, obtained at a temperature of 25 °C, for 100 µM WT Z7 in 100 mM HEPES with 50 mM NaCl, pH 7.5.

Figure 6C shows the increase in the initial velocity for hydrolysis as a function of the 4-NA substrate concentration. The slope of the plot gives a second order rate constant of 0.65 ± 0.03 M^-1^ s^-1^, a value comparable to the range between 0.23 and 0.57 M^-1^ s^-1^ reported for other tridentate catalytic ZNFs (Besold *et al*. 2016). Whereas the previously reported examples of catalytic ZNFs were designed polypeptides (Besold *et al*. 2016; Negi *et al*. 2004; Nomura and Sugiura 2004a, 2004b), Z7 may be the first example of a naturally occurring ZNF domain with hydrolytic activity.

### 2.7 WT Z7 has restricted aromatic ring rotation, and all three variants share thermal stability ≥ 80 °C

Some of the earliest evidence for protein dynamics came from observations of rapid ‘ring-flips’ that led to magnetic equivalence of protons on opposite sides of symmetric aromatic groups in proteins such as BPTI (Wuthrich and Wagner 1975) and HEWL (Campbell *et al*. 1975). Most tyrosine and phenylalanine aromatic rings in proteins undergo such ring-flips, characterized by 180° rotations about the Cβ-Cγ bond that results in degenerate NMR chemical shifts for the aromatic Hε1/Hε2 and Hδ1/Hδ2 protons. Ring flips are facilitated by protein ‘breathing motions’(Marino Perez *et al*. 2022) or ‘protein quakes’ that allow for aromatic ring rotation. It is rare to observe magnetically distinct signals for the protons on an aromatic ring (e.g. the five signals Hδ1, Hδ2, Hε1, Hε2, Hζ for a phenylalanine) since this requires restricted rotation that is typically associated only with very stable and rigid proteins. To our knowledge in ZNFs, restricted aromatic ring rotation has previously only been observed in a designed construct (Horx and Geyer 2021), and possibly the onset of restricted rotation was detected in one of the earliest studied naturally-occurring examples, Xfin-31 (Palmer *et al*. 1993).

In the TOCSY spectra of WT Z7 (Fig. 7A) we observe aromatic NMR spin systems comprised of five distinct signals for both F10 and F23. For the remaining aromatic group of Y1 we assigned a single crosspeak to a Hδ_1,2_-Hε_1,2_ correlation, however, there may be additional features in the spectrum that we could not resolve due to spectral crowding, so that Y1 may also be subject to restricted rotation. The non-degenerate signals from the WT Z7 aromatic groups of F10 and F23 show temperature dependences (Fig. 8A) typical of aromatic rings undergoing restricted 180° ring flips (Wuthrich and Wagner 1975, 1978). At low temperatures, the equivalent aromatic protons are in slow exchange on the NMR timescale. As the temperature is raised and protein dynamics are increased, the aromatic resonances broaden and coalesce as they enter the NMR intermediate exchange regime. At the highest temperatures, a single averaged chemical shift is expected for the magnetically equivalent resonances, however, in our case we were only able to observe an averaged signal for the F10 ε1/ε2 resonance due to spectral crowding.

**Figure 7.**
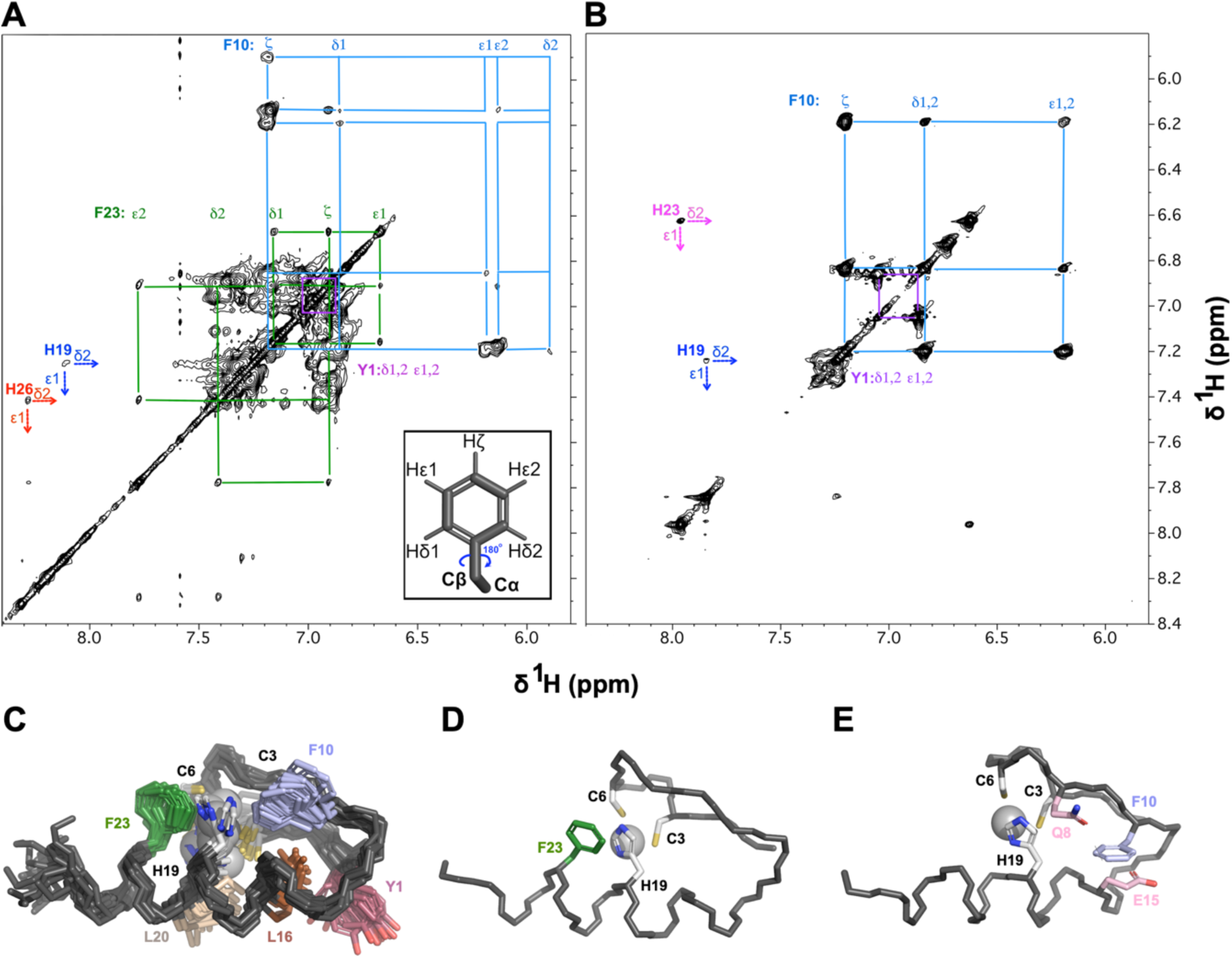
Aromatic regions of 70 ms mixing time 2D TOCSY spectra for samples in D_2_O. **(A)** WT. **(B)** Mut2. NMR spectra were obtained at 600 MHz. All aromatic spin systems in the two variants are indicated, except for H5 for which the Hε1-Hδ2 crosspeak is too weak in both spectra to be observed. The inset in A shows the aromatic ring of a phenylalanine, illustrating how 180° ring flips about the C2 axis delineated by the Cβ-Cγ bond, lead to magnetic equivalence of the δ and ε protons on opposite sides of the aromatic ring. This is the case in Mut2 (B) where degenerate NMR signals are observed for the Hδ1,2 and Hε1,2 protons of F23 indicating the phenylalanine is undergoing rapid ring-flips. F23 is not present in the Mut 2 sequence. In the WT (A), five non-degenerate aromatic proton signals are seen for each F10 and F23, indicating hindered rotation for both phenylalanines. **(C)** Illustration of hydrophobic sidechains and Zn^2+^ ligands in the NMR ensemble of WT Z7. The Zn^2+^ atoms are shown as spheres and the sidechains of the three ligands C3, C6, and H19 are shown in light gray. Hydrophobic residues Y1, F10, L16, L20, and F23 are in color. **(D)** NMR structure closest to the ensemble average illustrating how rotation of F23 is hindered by steric clashes with H19 in the Zn^2+^ coordination site. **(E)** Rotation of F10 is restricted due to residues Q8 and E18 (pink).

**Figure 8.**
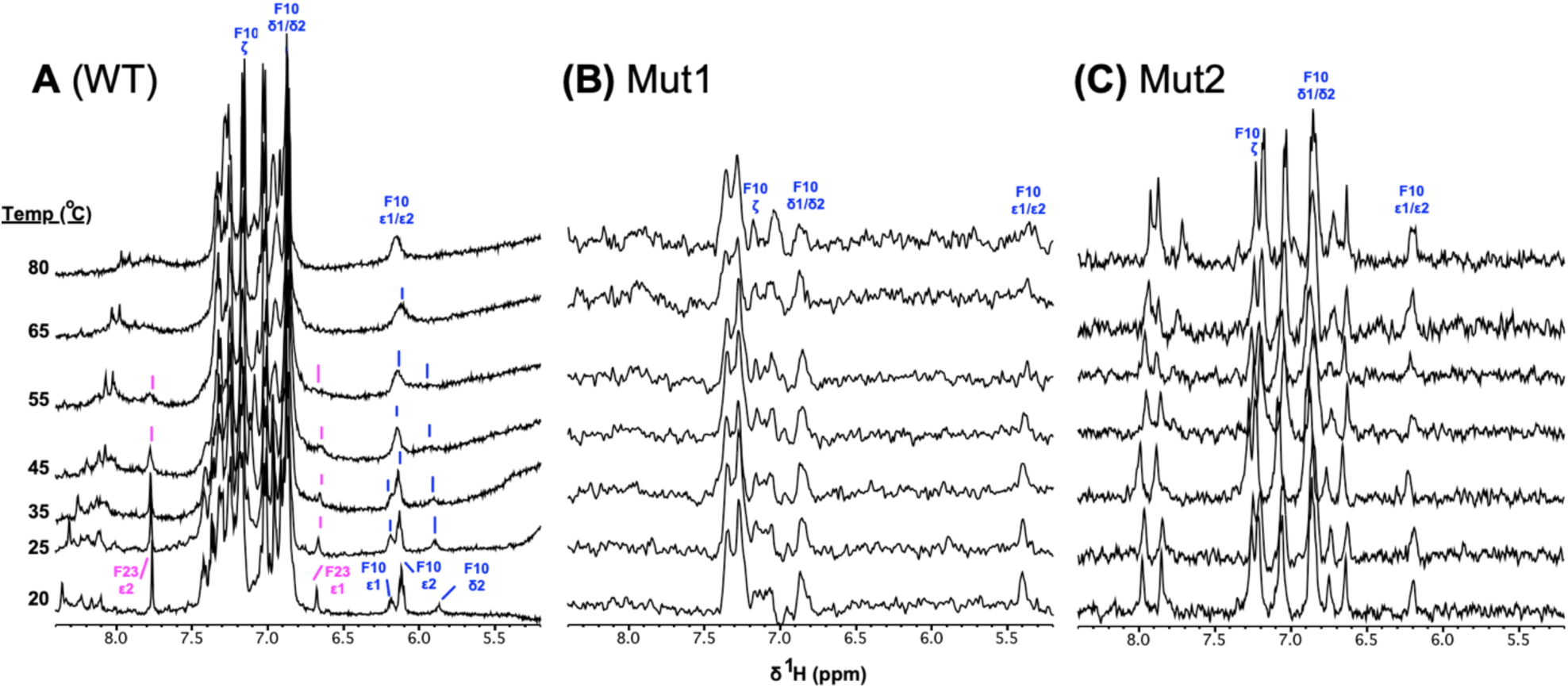
Temperature titrations monitored by 1D ^1^H NMR spectra. Aromatic regions of the NMR spectra of: **(A)** WT. **(B)** Mut1. **(C)** Mut2. All data were obtained at 500 MHz on samples dissolved in D_2_O, pH 6. For WT, the resolved ε1 and ε2 resonances of F23 as well as ε1 and ε2 resonances of F10 are in slow exchange at 20 °C. As the temperature increases, the rate of ring flipping moves from the slow to intermediate NMR exchange regimes, leading to exchange broadening of the NMR signals (indicated by the vertical lines). At 80 °C the ε1/ε2 resonances of F10 give an averaged shift corresponding to the fast exchange limit, while the ε1/ε2 resonances of F23 are not resolved in the 1D NMR spectra. For Mut1 and Mut2 the aromatic resonances of F10 give averaged signals consistent with the fast aromatic ring rotations throughout the entire temperature range.

In contrast to the WT ZNF, the TOCSY spectrum of Mut2 is markedly simplified (Fig. 7B). Mut2 has F23 missing due to the F23H substitution and both F10 and Y1 show simplified spin systems typical of aromatic groups undergoing rapid ring-flips. The poor solubility of the Mut1 mutant precluded 2D TOCSY experiments but 1D NMR spectra of Mut1 show a simplified aromatic region typical of rapid aromatic ring rotation (Fig. 8B). Figure 7C shows the aromatic and hydrophobic sidechains in the WT Z7 structure, which constitute a well-defined and closely packed nonpolar core for the domain. Based on the Z7 NMR structure, rotation of F23 is hindered by H19 which is bonded to the Zn^2+^ atom (Fig. 7D). Rotation of F10 appears to be obstructed by the polar residues Q8 and E15 (Fig. 7E). The switch to fast aromatic ring rotation in Mut1 and Mut2 probably does not involve rearrangements of the average structure compared to WT but rather an increase in protein breathing dynamics that allow the aromatic residues to rotate more freely. The temperature data in Figure 8 shows that all three variants remain folded up to a temperature of at least 80 °C, close to the limit of our experimental setup. While all three variants are exceptionally stable, the WT has reduced protein dynamics as evinced by the hindered rotation of its aromatic groups.

## 3. DISCUSSION

The folding status of the Z7 domain of ZNF711 has been an open question since the discovery of the gene encoding the transcription factor, when it was pointed out that the domain is unlikely to be functional because the position corresponding to the last histidine Zn^2+^ ligand is replaced by a phenylalanine (Lloyd *et al*. 1991). The UniProt database annotates the Z7 domain as degenerate at the time of this writing. The present work shows that the ZNF711-Z7 domain is a genuine zinc-finger that folds into a highly stable structure in the presence of Zn^2+^ (Fig. 1C, 2D). There is an increasingly prevalent view in modern biology that protein structures can be largely predicted by homology to the large number of known structures. The mis-annotation of the Z7 domain described in this work, our recent correction of the degenerate annotation for the Z* domain of Z750 (Rua *et al*. 2023), as well as several annotations we corrected previously (Rua *et al*. 2023) or are currently working on correcting, suggest there is still much to learn about the biophysical and structural properties of the resilient and versatile zinc-finger fold.

We initially thought that H26 could serve as the final Zn^2+^ ligand in the Z7 domain. The pH titration data indicate that this residue becomes protonated with a typical p*K*_a_ for a histidine (Fig. 3A,D) and is therefore not bonded to Zn^2+^. Moreover, the Mut1 variant with the H26A substitution retains the ability to fold in the presence of Zn^2+^, indicating the last histidine is not necessary for Zn^2+^-binding and folding (Fig. 1C). The data indicate that the WT Z7 domain has a tridentate binding site for Zn^2+^ comprised of just the three residues C3, C6, and H19. Precedent for tridentate ZNFs exists from protein design experiments and from disease-linked mutations that replace one of the Zn^2+^-ligating amino acids in the naturally occurring protein NEMO (Cordier *et al*. 2008). Cordier and Agou showed that the disease-linked C417F mutation removing the last ligand of a CCHC-type ZNF does not perturb the average three-dimensional NMR structure of the domain but increases its dynamics (Cordier *et al*. 2008). In early work, Berg’s group showed that deletion of the last four residues from the designed consensus ZNF CP-1, including the last metal-ligating H24, did not abrogate the ability of the peptide to bind Co^2+^ (Merkle *et al*. 1991). Nomura and Sugiura undertook a systematic substitution analysis of the four Zn^2+^-ligands in the CCHH-type second ZNF of Sp1 by glycine or alanine. They found mutants of each of the four chelating residues retained the ability to bind metal and had folded secondary structure by CD (Negi *et al*. 2004; Nomura and Sugiura 2002). Krezel’s group studied nine naturally occurring ZNFs with a tridentate XCHH, CXHH, CCXH, or CCHX ligand set rather than the tetradentate CCHH arrangement. All the tridentate ZNFs bound Co^2+^ and Zn^2+^, and metal-binding induced secondary structure by CD. The XCHH and CXHH types had lower metal affinities, the CCXH types tended to form ZnL_2_ complexes, while the CCHX-types had similar metal affinities and stabilities to the classical CCHH tetradentate ZNFs (Kluska *et al*. 2018a). Mackay’s group showed that mutation of the final histidine in the BKLF-F3 ZNF had indistinguishable effects on Zn^2+^-binding and did not affect the capacity of the domain to participate in functional DNA binding, concluding that the final zinc-chelating histidine is a non-essential feature of classical CCHH ZNFs (Simpson *et al*. 2003).

Our present work on the ZNF711-7 shows that tridentate ZNFs missing the last metal ligand can exist as naturally occurring wild type sequences with a stable well-defined tertiary structure as determined by NMR. One implication of this result is that the commonly used molecular biology approach of disrupting ZNF domain function by mutating Zn^2+^-chelating residues to alanine may be unsuitable (Simpson *et al*. 2003), particularly for the last ligand as the numerous examples of CCHX ligands given above demonstrate these can have normal ZNF function. The presence of tridentate CCHX ZNFs have the potential to confound current identification and classification schemes, illustrated by the fact that the UniProt database currently annotates the Z7 domain as degenerate. As described above, the occurrence of the functional tridentate Z7 domain was foreshadowed by protein design and protein engineering experiments (Merkle *et al*. 1991; Negi *et al*. 2023; Nomura and Sugiura 2002; Simpson *et al*. 2003). A second ZNF domain that occurs ZFX/ZFY, Z3, is thought to be inactive in ZNF711 since it is missing all the amino acids required to bind Zn^2+^ (Lloyd *et al*. 1991; Ni *et al*. 2020). Yet early protein design experiments from the Imperiali group showed that it is possible to create a zinc-less stably-folded ZNF buttressed by hydrophobic interactions in lieu of a metal binding site (Struthers *et al*. 1996). Whether the segment corresponding to the Z3 domain of ZFX/ZFY is unfolded or plays a structural role in ZNF711 is an open question that will ultimately need to be addressed experimentally.

The putative H26 ligand does not participate in metal coordination in the tridentate Z7 domain. In contrast, a histidine introduced at the consensus position through the F23H substitution in the Mut2 variant (Fig. 1B), bonds to Zn^2+^ based on pH titrations experiments (Fig. 3C,F) and gives an increased d-d absorption band characteristic of tetrahedral coordination in UV-Vis spectra of the Co^2+^ substituted polypeptide (Fig. 5B). The preference for a histidine to coordinate Zn^2+^ at position 23 but not at position 26 appears to be dictated by the location of the potential ligands in the structure. An X3 spacing between the third and fourth Zn^2+^ ligands is strictly conserved in all odd ZNF domains of the ZFX/ZFY/ZNF711 family (Lloyd *et al*. 1991), whereas X4 is conserved for the even-numbered domains. An X3-X5 spacing between the last two ligands is preferred by ZNFs in general (Rua *et al*. 2023). Coordination by H26 would need an X6 spacing. An early protein engineering experiment from the Berg lab showed that most residues in a ZNF could be replaced by alanines and the ZNF still folded, as long as the ligand spacing and the conserved hydrophobics were retained (Michael *et al*. 1992). Although H26 does not participate in Zn^2+^ binding, and chemical shift-derived *S*^2^ order parameters suggest the residue is flexible, the H26A mutation is not innocuous. The Mut1 variant harboring the H26A mutation has a lowered NMR chemical shift dispersion, and a CD spectrum closer to the unfolded domain than that of Zn^2+^-bound WT. Both the Mut1 and Mut2 variants have unrestricted aromatic residue rotation in contrast to the WT Z7 domain.

The *K*_d_ values of the tridentate WT and Mut1 domains for Co^2+^ and Zn^2+^ are similar to those of the tetradentate Mut2 variant (Table 2), indicating there is no direct relationship between the avidity of the Z7 domain for metals and the number of ligands employed in chelation. While the WT domain gives the lowest *K*_d_ values with both metals, those for the Mut1 and Mut2 mutants are within a factor of two for Co^2+^ and five for Zn^2+^. All three variants are extremely stable with thermal melting points above 80 °C (Fig. 8). That there are no large differences in metal binding or stability between these variants begs the question of why the vast majority of ZNFs have four metal-ligating residues rather than three. Moreover, a recent survey found that the most common cancer mutations that occur within ZNF domains, involve the substitution of the last metal-ligating residue (Munro *et al*. 2018). Perhaps these cancer mutations do not abrogate Zn^2+^-binding but introduce harmful new function – a hypothesis that warrants further investigation. Like two previously described ZNFs with only three ligands (Besold *et al*. 2016; Nomura and Sugiura 2004a), the WT Z7 domain has hydrolytic activity towards the substrate 4-NA (Fig. 6). This type of catalytic activity has the potential to turn a structural or DNA-binding ZNF module into one that can cleave DNA, although to date this has only been demonstrated for an engineered tandem of three HHHH ZNFs and not yet for a CCHX type ZNF (Negi *et al*. 2023; Nomura and Sugiura 2004b). While the acquisition of hydrolytic activity through the mutation of the last metal ligand would be detrimental, it might be tolerated for Z7 if the domain is shielded in the overall structure of the ZNF711 transcription factor away from the DNA binding site. The evolutionary driver for the loss of the last ligand in Z7 may have been tighter metal binding, as demonstrated by the tridentate WT domain having a lower *K*_d_ for Zn^2+^than the tetradentate Mut2 mutant, albeit with only a factor of five difference.

Transcription factors are considered notoriously undruggable due to their lack of defined drug binding sites (Tao and Wu 2023). For ZNF-containing transcription factors some success has been achieved using ‘zinc-ejectors’ -compounds that form irreversible dead-end complexes with the Zn^2+^-binding ligands in ZNFs (Brue *et al*. 2022; Harney *et al*. 2009; Rice *et al*. 1993). It would be interesting to see if tridentate ZNFs such as Z7 are more susceptible to Zn-ejectors than their more common tetradentate counterparts. Any strategies to improve the druggability of ZNFs must deal with the high repetition of these domains in ZNF-array containing proteins, and the high homology between transcription factor paralogs. If zinc-ejectors are more effective with domains having unusual tridentate metal ligating sites like that of Z7, this could provide an extremely useful therapeutic handle to (i) target the exceptional domain without affecting the others in a ZNF-array protein, and (ii) specifically target a transcription factor like ZNF711 involved in the genetic XLID97 disease without affecting the highly homologous ZFX and ZFY paralogs (Johnston *et al*. 1998; Lloyd *et al*. 1991).

## 4. MATERIALS AND METHODS

### 4.1 Materials

ZnSO_4_•H_2_O (purity ≥ 99.9%) was from Sigma (St. Louis, MO). 99.96% D_2_O was from Cambridge Isotopes (Tewksbury, MA), and 4-nintophenyl acetate (4-NA) was from Sigma (St Louis, MO). The 27-residue Z7 peptide with the WT sequence, corresponding to the fragment 562-588 of human ZNF711 (UniProt Q9Y462), was custom synthesized at 96% HPLC purity by Biomatik (Kitchener, Canada). The Mut1 peptide was synthesized at 96% purity by AAPPTec (Louisville, KY) and Mut2 was made at 86% purity by Biomatik. All three peptides had their termini blocked by N-acetylation and C-amidation and had molecular weights by mass spectrometry within 1 Dalton of those theoretically expected from the sequence. Peptide concentrations were determined by absorption at 280 nm (A_280_) using a molar extinction coefficient of 1,559 M^-1^ cm^-1^ for the unfolded Z7 in the absence of Zn^2+^. The peptide concentration to determine the extinction coefficient was determined with a BCA assay (Walker 2002). The value for the unfolded peptide is very close to that of 1,490 M^-1^ cm^-1^ predicted from its sequence with the ProtParam program (Gasteiger *et al*. 2005). For folded ZNF711 in the presence of Zn^2+^, we determined an A_280_ extinction coefficient of 2,383 M^-1^ cm^-1^.

### 4.2 CD measurements and Zn^2+^ affinity determination

Circular dichroism (CD) data were collected using an Applied Photophysics Chirascan V100 Spectrometer (Surrey, UK) using a 1 mm cuvette. Z7 peptide concentrations were 51 µM for WT and 75 µM for Mut1 and Mut2, in 10 mM NaPO_4_ buffer, pH 7. CD spectra were collected between 190-250 nm, using a 1 nm bandwidth, a 1 nm step size, and 5 s/point data averaging, for an approximate total scan time of 5 minutes. Zn^2+^-binding was measured in a competition assay with 17.71 mM EGTA (Rua *et al*. 2023), with 0.2 mM TCEP (tris(2-carboxyethyl)phosphine) included as a reducing agent to prevent cysteine disulfide formation. Because the affinity for Zn^2+^ is highly depended on pH, WT and mutant samples were carefully maintained at a constant pH of 7.0 for all metal-binding experiments. *K*_d_ values were calculated from the EGTΑ competition experiments as previously described (Rua *et al*. 2023).

### 4.3 UV-Vis characterization of Co^2+^ binding

Co^2+^ binding experiments were performed on an Ultrospec 8000 UV-Vis spectrophotometer (Thermo Fisher) for 100 µM Z7 peptide samples in 10 mM Tris pH 7.0, containing 0.5 mM of the reducing agent TCEP. Stock solutions of 5, 40, and 400 mM CoCl_2_ were used to titrate samples to a range of Co^2+^ concentrations between 0 to 300 µM. The increase in A_630 nm_ as a function of Co^2+^ concentration was fitted to a sigmoidal function to obtain *K*_d_ (Rua *et al*. 2023).

### 4.4 Hydrolytic activity with the 4-nitrophenyl acetate (4-NA) substrate

The hydrolysis of 4-NA to 4-nitrophenolate (4-NP) and acetate (Fig. 6A) was monitored by A_400nm_ using an Ultrospec 8000 UV-Vis spectrophotometer. Experiments were done at 25 °C, for samples in 100 mM HEPES buffer pH 7.5, 50 mM NaCl. The 4-NA was dissolved in acetonitrile and added to the samples to a final concentration of 1% (v/v) acetronitrile. The Z7 peptide with the WT sequence was 100 µM with an equimolar concentration of ZnSO_4_. To measure kinetics, concentrations of 4-NA ranging from 0-200 μM were added, and A_400nm_ was sampled using a path length of 1 cm, every 10 s for 1000 s. Initial rates were obtained using an extinction coefficient of 12,800 M^-1^cm^-1^ for the reaction product 4-NP at pH 7.5 (Nomura and Sugiura 2004a). The second order rate constant was calculated as described in the literature (Besold *et al*. 2016).

### 4.5 NMR spectroscopy

Since the peptides were purified as TFA salts, freshly dissolved samples for NMR typically had pH values between 2 and 3. Under these conditions the peptides are unfolded and readily soluble to concentrations up to 12 mM. To form the Zn^2+^-bound peptides, ZnSO_4_ dissolved as appropriate in H_2_O or D_2_O was added in the desired amount to the peptides at acidic pH, and the pH was adjusted with NaOH/HCl or NaOD/DCl. As the pH was raised, some precipitation occurred near the isoelectric points (pIs = 6.0 to 6.3) of the WT and Mut1 peptides. Precipitation was more severe for Mut1 than WT and was observed at NMR concentrations above 1 mM, but not at the lower peptide concentrations used for CD and UV-Vis spectroscopy. Precipitates were removed by sedimentation. Pulse-field gradient diffusion experiments (Sallum *et al*. 2007; Whitehead *et al*. 2022) calibrated against the internal standard DSS, gave an R_h_ of 9.1 ± 0.6 Å for a Zn^2+^-bound 2.4 mM WT sample at 25 °C, pH 6.2, 600 MHz, consistent with a monomeric oligomerization state for the Z7 domain under NMR conditions (Rua *et al*. 2023; Whitehead *et al*. 2022).

Temperature titrations for Zn^2+^-bound WT, Mut1, and Mut2 were done in D_2_O at pH 6.2, on a 500 MHz Bruker Avance instrument with a room temperature probe. After each temperature ramp from 20 to 80 °C, an NMR spectrum was obtained at 25 °C after cooling from 80 °C, as a reversibility check to confirm that the samples were not damaged by heating.

pH titrations were done on a Varian INOVA 600 MHz instrument with a cryogenic probe at a temperature of 25 °C using samples dissolved in D_2_O. Solution pH values were measured with a MA235 pH meter equipped with a glass InLab Micro pH electrode from Mettler (Columbus, OH). The pH value was taken as the average of measurements before and after NMR experiments. The pH values were not corrected for the deuterium isotope effect, since the isotope effect on pH is comparable and opposite in magnitude to the isotope effect on a glass pH electrode (Croke *et al*. 2011; Kaplan *et al*. 2017). For titrating histidines, p*K*_a_ values were calculated from fits of the data to a Henderson-Hasselbalch equation as previously described (Harprecht *et al*. 2016).

### 4.6 NMR assignments

Experiments for NMR assignments were done at a temperature of 25 °C on a Bruker Avance 600 MHz instrument equipped with a TCI cryogenic probe that was part of the Francis Bitter Magnet Lab of MIT. A sample containing 2.4 mM of the WT Z7 peptide (determined by A_280_) and 4.4 mM ZnSO_4_ in 90% H_2_O/10%D_2_O at pH 6.2, was used to collect 2D TOCSY (70 ms mix time), NOESY (200 ms mix time), DQF-COSY, and natural abundance ^15^N-sofast-HMQC spectra. The sample was lyophilized and resuspended in an equivalent volume of 99.96% D_2_O to identify exchange-protected amide protons using 1D ^1^H-NMR spectra. A 2D TOCSY experiment on the D_2_O sample was collected to assign aromatic spin systems, and a ^1^H-^13^C HSQC spectrum was used for ^13^C assignments. NOESY (50 ms mix time) and E.COSY spectra for the D_2_O sample were used for stereospecific methylene proton assignments (Case *et al*. 1994). The extent of assigned resonances was ^1^H (93%), backbone ^15^N (92%) and ^13^C (67%, excluding C’). The ^15^N assignments should be considered tentative, however, since the ^15^N-sofast-HMQC experiment had poor sensitivity at natural isotope abundance (0.36% for ^15^N). ^1^H chemical shifts were referenced to internal DSS (2,2-dimethyl-2-silapentane-5-sulfonate), while ^13^C and ^15^N shifts were referenced indirectly as described in the literature (Wishart *et al*. 1995).

### 4.7 NMR Structure calculations

Distance restraints were obtained from a 250 ms NOESY spectrum collected at a temperature of 10 °C on an 800 MHz Bruker instrument equipped with a cryoprobe. While the data for NMR assignments was recorded at 25 °C, the HN proton of L16 is barely visible at this temperature (inset Fig 1C) but becomes stronger at 10 °C facilitating assignments and structure determination (Fig. 2B). The remainder of the ^1^H assignments were conserved and transferrable from 25 °C to 10 °C. Torsional restraints were calculated from ^1^H_N_, ^1^Hα ^13^Cα, ^13^Cβ, and ^15^N chemical shifts using the program TALOS-N (Shen and Bax 2015). The three Zn^2+^-ligating residues were identified from pH titration experiments as described in the Results section. Distance restraints from C3 and C6 to the Zn^2+^ atom, were set to 2.33-2.37 Å for Zn^2+^-Sγ and 3.25-3.51 Å for Zn^2+^-Cβ bonds, respectively. An ambiguous restraint of 1.0-3.1 Å was set between Zn^2+^ and either the Nδ1 or Nε2 atom of H19. Hydrogen bond restraints were included for amide protons determined to be in hydrogen-bonded secondary structure (Fig. 2C). Two distance restraints were included per H-bond (1.5-2.5 Å for NH-O and 2.5-3.5 Å for N-O) to enforce linearity. An initial set of 100 structures were calculated with the program XPLOR-NIH (Schwieters *et al*. 2003) on the NMRbox platform (Maciejewski *et al*. 2017), from which the 20 lowest energy structures without violations were selected for deposition and analysis.

### 4.8 Database accession numbers

NMR assignments and chemical shift values for the Z7 domain were deposited in the BMRB under accession code 52210. Coordinates and restraints for NMR structure calculations, were deposited in the PDB under accession code 8VG3.

## AUTHOR CONTRIBUTIONS

**A.J. Rua:** Investigation; writing – review and editing; **A.T. Alexandrescu:** Conceptualization; investigation; writing – original draft, review and editing; visualization; resources; supervision.

## ACKNOWLEDGEMENTS

We thank Prof. Carolyn Teschke’s for use of her group’s CD and UV-Vis spectrophotometers. We thank Prof Mei Hong for use of her group’s solution 600 MHz instrument during the sabbatical of A.T.A. The NMR experiments used equipment at the MIT-Harvard Center for Magnetic Resonance, which is supported by the P41 grant GM132079. M.H. is partially supported by NIH grant AG059661. We acknowledge use of the NMRbox platform (https://nmrbox.nmrhub.org) for NMR data analysis and structure calculation.

